# Linking heartbeats with the cortical network dynamics involved in self-social touch distinction

**DOI:** 10.1101/2024.05.15.594340

**Authors:** Diego Candia-Rivera, Fabrizio de Vico Fallani, Rebecca Boehme, Paula C. Salamone

## Abstract

Research on interoception has revealed the role of heartbeats in shaping our perceptual awareness and embodying a first-person perspective. These heartbeat dynamics exhibit distinct responses to various types of touch. We advanced that those dynamics are directly associated to the brain activity that allows self-other distinction. In our study encompassing self and social touch, we employed a method to quantify the distinct couplings of temporal patterns in cardiac sympathetic and parasympathetic activities with brain connectivity. Our findings revealed that social touch led to an increase in the coupling between frontoparietal networks and parasympathetic/vagal activity, particularly in alpha and gamma bands. Conversely, as social touch progressed, we observed a decrease in the coupling between brain networks and sympathetic dynamics across a broad frequency range. These results show how heartbeat dynamics are intertwined with brain organization and provide fresh evidence on the neurophysiological mechanisms of self-social touch distinction.

## Introduction

The feeling that your body is yours involves the integration of multiple sensory inputs^1^, including the sensing, processing, and representation of bodily signals, typically referred to as interoception^2^. This meta-representation of bodily signals allows to define the bodily self, i.e., to define which bodily parts are yours^3^ and to establish the boundaries between the self, others, and the environment^4,5^. However, it remains unclear to what extent interoceptive inputs contribute to the neural processes underpinning the self-other distinction. One of the proposed mechanisms contributing to these distinctions is the prediction, and posterior suppression, of the sensory effects of one’s own actions^6^, but a longstanding discussion remains about how the brain uses that efferent information^7^.

Touch is considered an essential contributor to the establishment and maintenance of the bodily self^7^. Touch interactions involve both exteroceptive processing, which is the perception of tactile information, and interoceptive processing, which includes the processing of resulting physiological changes such as heart rate fluctuations^2^. Experimental evidence suggests that the neural formation of the bodily self involves the interplay of multiple cortical regions. For instance, the premotor cortex has been associated with the feeling of body ownership^8,9^, the somatosensory cortex with the attribution of seen touch to felt touch^10^, the insular and anterior cingulate cortices with social touch^11,12^, the posterior superior temporal sulcus with more general social cognition processing^13^, and the right superior temporal lobe with the conscious processes that allow tracking self-generated actions^14^. Complementarily, distinct activations in the spine reflect the differences between self-generated touch and social touch^12^. Indeed, these mechanisms are disrupted after spinal cord injury, and hypothesized that it is caused from the loss of sensory and motor functions^15^, which may cause an a posteriori disruption in affective and social touch processing/perception^16^.

In this study, we focus on skin-to-skin touch performed at an optimal stroking speed that activates C-fibers, resulting in a pleasant sensation and often referred as affective touch^17,18^. Research employing EEG/MEG under different touch paradigms has unveiled insights for various frequency bands. Touch modulates a wide EEG spectrum within theta-gamma oscillations, with alpha and beta bands mostly reported^19–25^. While the exact specificity of these EEG responses is still uncertain, they have been associated with both sensory processing and emotional regulation^20,22^. For instance, while beta connectivity has been suggested as a mechanism for the processing of somatosensory stimuli, alpha connectivity presents some specificity for conscious somatosensory processing^26,27^. Exploring brain oscillations during touch could enhance our understanding of its physiological basis. However, there is limited evidence connecting widespread interactions between brain and peripheral neural dynamics, encompassing interoceptive and regulatory mechanisms.

One of the proposed mechanisms to differentiate between self- and social touch involves the integration and matching of information coming from tactile and proprioceptive inputs^5^. While interoception has been proposed as a relevant factor in affective touch^2,28–30^, there is limited understanding of the role interoceptive inputs play in distinguishing social touch compared to self-produced touch. In that line, we recently showed that heartbeat dynamics contribute to the distinction of social and self-touch^31^. However, studies on touch and its associated neural pathways often neglect to consider potential connections with the autonomic system, including vagal pathways^32^. Furthermore, disruptions in the autonomic system that may affect temperature perception or pain sensitivity do not necessarily impact affective touch^33^, demonstrating the specificity of the pathways involved in this type of tactile experience. Therefore, in this study, we aimed to identify the brain dynamics specific to social touch. We were specifically interested in the difference between self-touch and social touch and hypothesized that the social component generates unique brain-heart interactions, encompassing frontoparietal connections^34–37^, which enable the neural distinction between self and others. To control whether the effects observed during self-touch were driven by the motor-component, we also compared self-touch to object-touch. We expected self-touch to differ from object-touch considering that it involves an additional sensory component (the touched arm) and probably a specific prediction-model underlying sensory attenuation observed on behavior and neural correlates during self-directed touch^12,38–40^.

Previous endeavors have linked interoceptive mechanisms with affective touch^2,41^, body-ownership^42^, perspective-taking^43–45^, and consciousness^46,47^. We proposed that interoceptive mechanisms play a key role in shaping the neural differences between social touch and self-generated touch. These mechanisms may be part of the integral components behind the intricate neural processes that contribute to our subjective and profound tactile experiences^48^. Additionally, we postulate that such mechanisms can be quantified through the analysis of brain-heart interactions. We tested a recently proposed framework to study brain-heart interplay by quantifying the relationship between brain connectivity and estimators of cardiac sympathetic and parasympathetic activities^49^. We aimed to determine the physiological changes triggered by the touch conditions as reflected in brain-heart interplay estimated from EEG and ECG data in a cohort of 28 healthy adults who underwent a multimodal touch paradigm^12^. Our findings uncover a prominent role played by the coupling between alpha and gamma brain connectivity, and parasympathetic/vagal activity in social touch, highlighting the role of interoceptive mechanisms in the context of self-other distinction, embodiment, and affective touch.

## Results

We studied brain-heart interactions in healthy participants undergoing a multimodal touch paradigm^12^. The protocol consisted in the recording of EEG and ECG data while undergoing three distinct conditions: social touch (being stroked on the forearm by the experimenter), self-touch (stroking of the participant’s own forearm), and object-touch (participant stroking a pillow) as a control condition. Each of the three conditions lasted 180 seconds, that were analyzed in the segments 0-60, 60-120 and 120-180 seconds, as done previously^31^.

In our previous results, HRV revealed notable differences in sympathetic and parasympathetic responses based on touch type and time interval^31^. For cardiac sympathetic indices, significant differences were observed between self- and social touch across all intervals, with social touch showing lower indices compared to self-touch. Social touch also had lower indices than object touch, though the difference between self- and object touch was not always significant. For cardiac parasympathetic indices, social touch consistently had higher values compared to self-touch. No significant differences were found between self- and object touch, but social touch had higher parasympathetic indices than object touch in all intervals (see Supplementary material, Tables S2 and S3).

In the analyses presented here, our physiological data analysis focused on identifying the distinct cortical networks that dynamically form in conjunction with the previously described fluctuations in cardiac dynamics, for each touch modality, at different frequency bands and latencies.

We used a method that quantifies the coupling between brain connectivity derived from EEG data, and cardiac sympathetic and parasympathetic activities obtained from ECG recordings^49^, which is represented in a general scheme in Figure 1. We compared brain-heart coupling matrices that depict the relationship between each pair of EEG channels in relation to cardiac dynamics across the different touch modalities of this study: social vs. self, social vs. object, and self vs. object touch.

**Figure 1.**
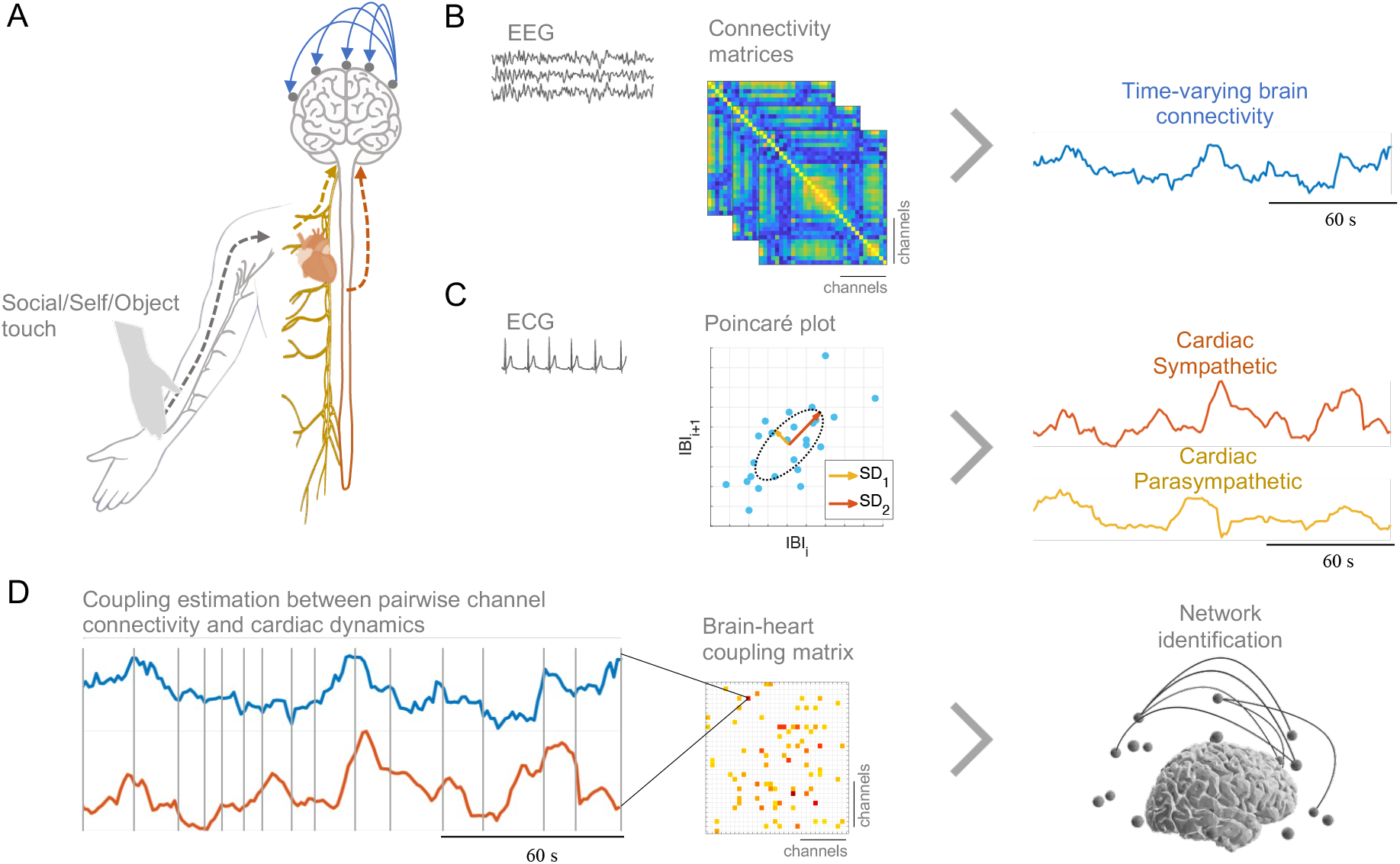
Brain connectivity-Cardiac Coupling under touch.(A) The framework aims to quantify the brain-heart coupling under three modalities of touch: social, self-, and object-touch. (B) The framework involved the computation of brain connectivity from EEG and (C) the estimation of cardiac sympathetic and parasympathetic/vagal activities are computed from the successive changes in interbeat intervals (IBI) gathered from the Poincaré plot (for an in depth description of this method see^50^). (D) The coupling quantification is achieved by assessing the similarities between two time series, using the Maximal Information Coefficient (MIC) method, which evaluates the similarities between distinct segments individually, using an adjusted grid as depicted in the figure. The overall measure combines the similarities observed throughout the entire time-course. From the brain-heart coupling matrices, the networks are identified by grouping neighboring links using a nonparametric permutation test.

The network size depicted in Figure 2 is defined as the number of connections linking two EEG channels within a specific frequency band, and that these connections changed their coupling with cardiac activities as a function of the touch modality. This network exhibits a specific coupling with either cardiac sympathetic or parasympathetic indices for the distinction of the touch modalities. The results indicate that the most pronounced distinctions are observed between social and self-touch, with prominent differences during the early stages (0-60 seconds). These distinctions primarily manifest in the coupling of parasympathetic activity with alpha, beta and gamma networks. Furthermore, in the late stages (120-180 seconds), these distinctions become more prominent in the coupling with sympathetic activity, where self-touch presents higher brain-heart coupling than social touch. Regarding the distinction between self and object touch, we found that brain-heart coupling is more influenced by parasympathetic activity and tends to be higher in later stages. A detailed statistical comparison of the self-vs social touch is presented in Table 1 (for comparisons with object-touch modality, see Supplementary material, Table S1).

**Table 1.** Results from the identification of brain networks whose couplings with cardiac sympathetic or parasympathetic activity differentiate between social vs self-touch. The statistical analyses are based on the cluster permutation test, where the cluster size indicates the number of connections whose coupling with cardiac dynamics distinguished social vs self-touch. These analyses were performed separately per EEG frequency band, and their couplings with either cardiac sympathetic or parasympathetic activities. The reported Z-values correspond to the range of Z-values within the cluster. P-values correspond to the cluster p-value obtained from 10,000 permutations.

| | | $\delta$ | $\theta$ | $\alpha$ | $\beta$ | $\gamma$ |
| --- | --- | --- | --- | --- | --- | --- |
| Brain-parasympathetic | 0-60s | size=6, Z=2.03–2.82, p=0.0007 | - | size=10, Z=2.03–2.53, p=0.0000 | size=12, Z=2.00–2.62, p=0.0005 | size=19, Z=2.00–2.66, p=0.0000 |
|  | 60-120s | - | - | size=4, Z=2.00–2.41, p=0.0021 | size=2, Z=2.44–2.46, p=0.0023 | size=2, Z=2.09–2.19, p=0.0070 |
|  | 120-180s | - | - | - | - | - |
| Brain-sympathetic | 0-60s | - | - | - | - | - |
|  | 60-120s | - | - | - | - | size=3, Z=2.05–2.48, p=0.0020 |
|  | 120-180s | size=33, Z=1.98–4.42, p<0.0001 | size=26, Z=1.98–3.12, p<0.0001 | size=21, Z=1.98–3.78, p<0.0001 | size=4, Z=2.03–2.71, p=0.0071 | size=39, Z=2.00–3.48, p<0.0001 |

**Figure 2.**
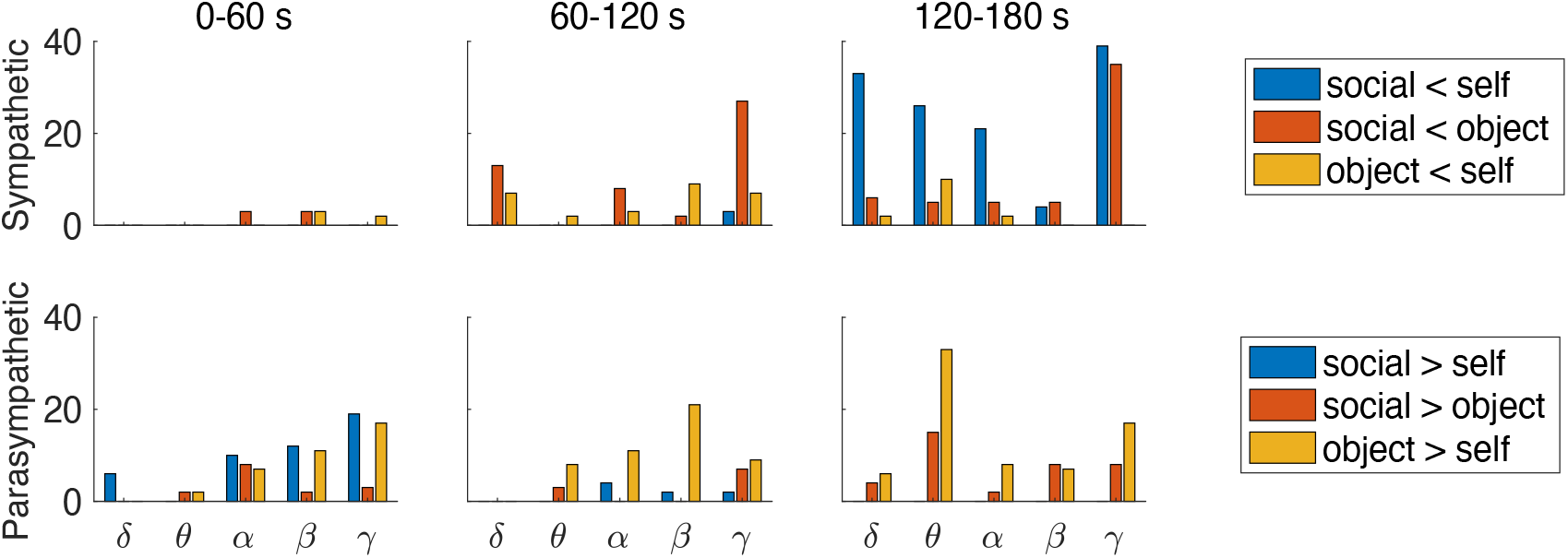
Summary of the results on the identification of brain networks whose couplings with cardiac sympathetic or parasympathetic activity differentiate between modalities of touch. The comparisons encompass social vs. self-touch, social vs. object-touch, and self vs. object touch. The statistical analyses are based on the cluster permutation test to identify the brain networks that showed an increased or decreased coupling with respect to the touch modalities compared, as specified in the legend. The histograms display the network size, which indicate the number of EEG connectivity links that presented a significant change in their coupling with cardiac activity, between the two touch modalities compared.

In the early stages of touch, there was an increase in brain-heart coupling specific to social touch, precisely between cardiac parasympathetic activity and a broad spectrum of EEG bands, when compared to self-touch. Those results suggest that the social component causes a rapid physiological modulation encompassing both brain and cardiac parasympathetic dynamics. In Figure 3, we illustrate the main networks that exhibited a distinct coupling with cardiac parasympathetic activity, distinguishing between social and self-touch within the 0-60 seconds interval. The alpha network connecting the frontal lobe to midline parietal regions, showed an elevated coupling with parasympathetic activity during social touch compared to self-touch. The beta network interconnecting the right central areas with both anterior and posterior regions, displayed an increased coupling with parasympathetic activity during social touch as opposed to self-touch as well. Lastly, the gamma network with higher coupling with parasympathetic activity during social touch, encompassed parietal to frontal connections.

**Figure 3.**
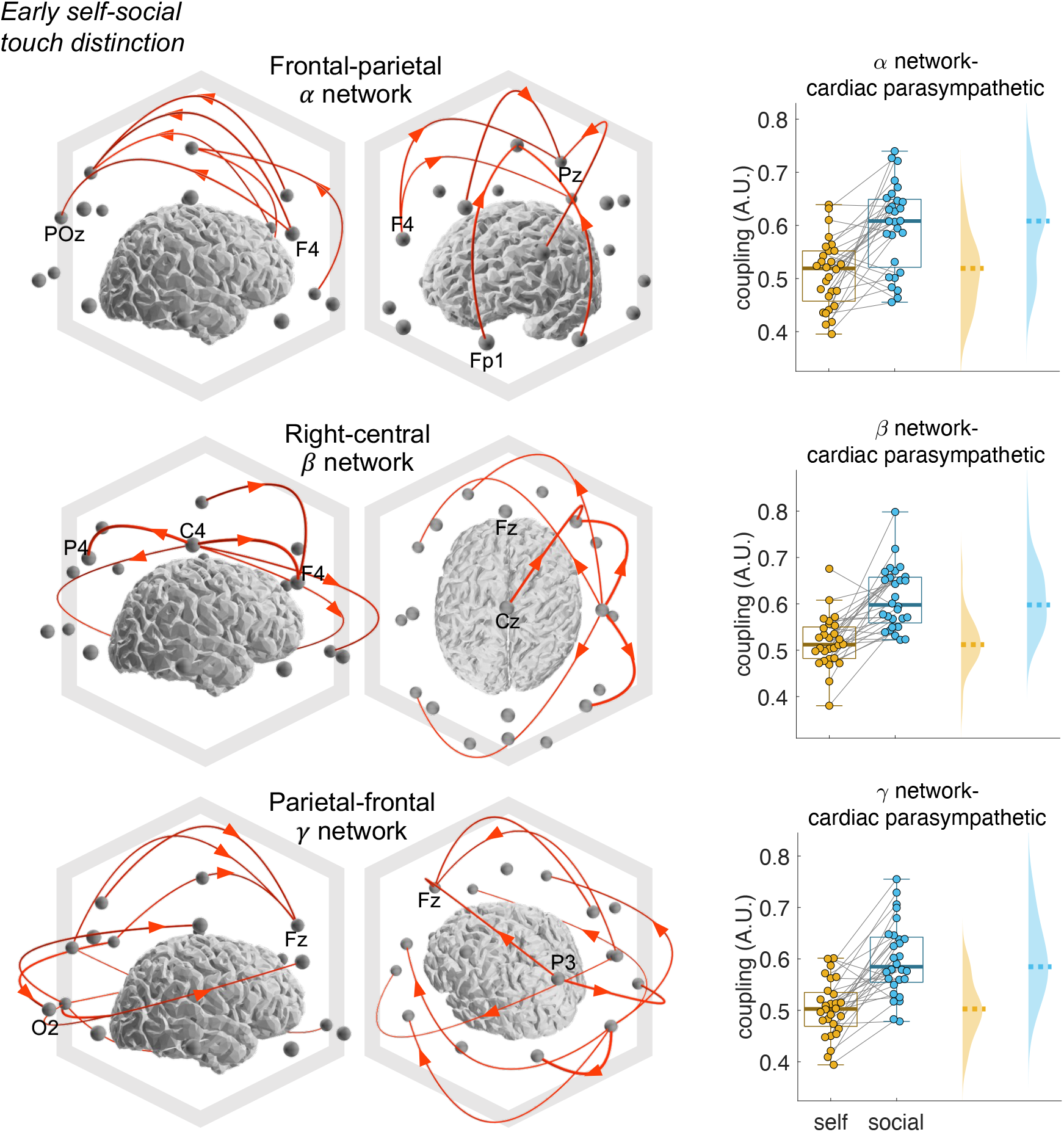
Brain-heart coupling distinguishing social vs self-touch at the early stage (0-60s). The main distinctions were found in the coupling between alpha, beta, and gamma brain networks with cardiac parasympathetic activity. Main brain networks were defined as network size > 10 connections. The arrows indicate the directed EEG connectivity links pertaining to the identified network. The right column displays the average coupling across all links pertaining to the identified network. Coupling ranges between 0-1, indicating the degree of co-fluctuations between brain and heart signals. Each data point corresponds to one participant.

Our findings indicate that the initially elevated brain-heart coupling observed during social touch, particularly with parasympathetic activity in comparison to self-touch, diminished after 60 seconds and returned to previous levels after 120 seconds. Intriguingly, later stages of social touch revealed a decreased brain-heart coupling specifically between cardiac sympathetic activity and a broad EEG connectivity spectrum. These results imply that prolonged social touch induces a strong physiological response encompassing global brain dynamics and its detachment with cardiac sympathetic inputs. In Figure 4, we depict the main networks that presented a difference in the coupling with cardiac sympathetic activity, distinguishing between social and self-touch within the 120-180 seconds interval. Within the delta band, we observed connections that exhibited reduced coupling with sympathetic activity during social touch in the frontal-parietal regions, in both directions. In the theta band, the primary network with reduced coupling with sympathetic activity is located in posterior regions under social touch. The alpha networks, which connect the frontal lobe to parietal regions, displayed reduced coupling with sympathetic activity during social touch compared to self-touch. Similarly, the gamma network exhibited lower coupling with sympathetic activity during social touch and featured frontal-to-parietal connections.

**Figure 4.**
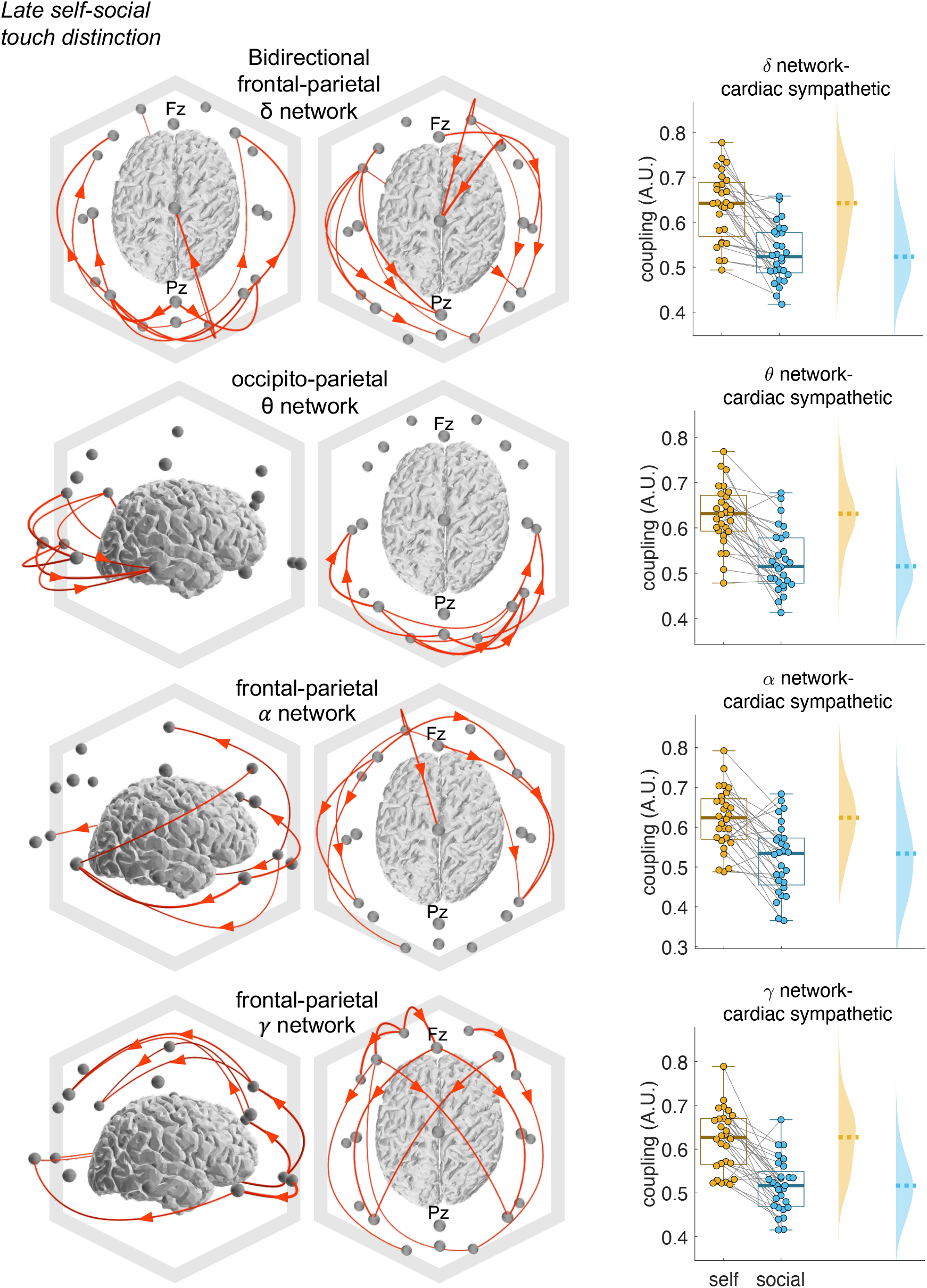
Brain-heart coupling distinguishing social vs self-touch at the late stage (120-180s). The main distinctions were found in the coupling between delta, theta, alpha, and gamma brain networks with cardiac sympathetic activity. Main brain networks were defined as network size > 10 connections. The arrows indicate the directed EEG connectivity links pertaining to the identified network. The right column displays the average coupling across all links pertaining to the identified network. Coupling ranges between 0-1, indicating the degree of co-fluctuations between brain and heart signals. Each data point corresponds to one participant.

Notably, when controlling self-social touch distinctions solely for changes in EEG connectivity, we observed parallel increases and decreases in EEG connectivity (Figure 5A), with an overall smaller network size as compared to the brain-heart coupling counterparts. Remarkably, we found that EEG connectivity patterns distinguishing self-social touch do not always follow the same direction as those observed in brain-heart coupling measures. In the late touch stage (3^rd^ minute), we observed simultaneous increases and decreases (Figure 5B and C). Moreover, those changes appear to involve fronto-parietal connections (Figure 5D), as seen in the brain-heart coupling analysis. These results suggest that social touch affects both HRV and EEG connectivity, but our study highlights a relationship between these parallel, physiological phenomena, suggesting that responses to touch involve interconnected brain-heart responses, rather than isolated mechanisms. Moreover, since the brain-heart coupling analysis involves a larger network compared to EEG connectivity alone, it appears that we are not just capturing increases or decreases in connectivity but rather the co-fluctuation of brain and cardiac dynamics. The degree of this coordination, and the structures involved, are significantly influenced by social touch.

**Figure 5.**
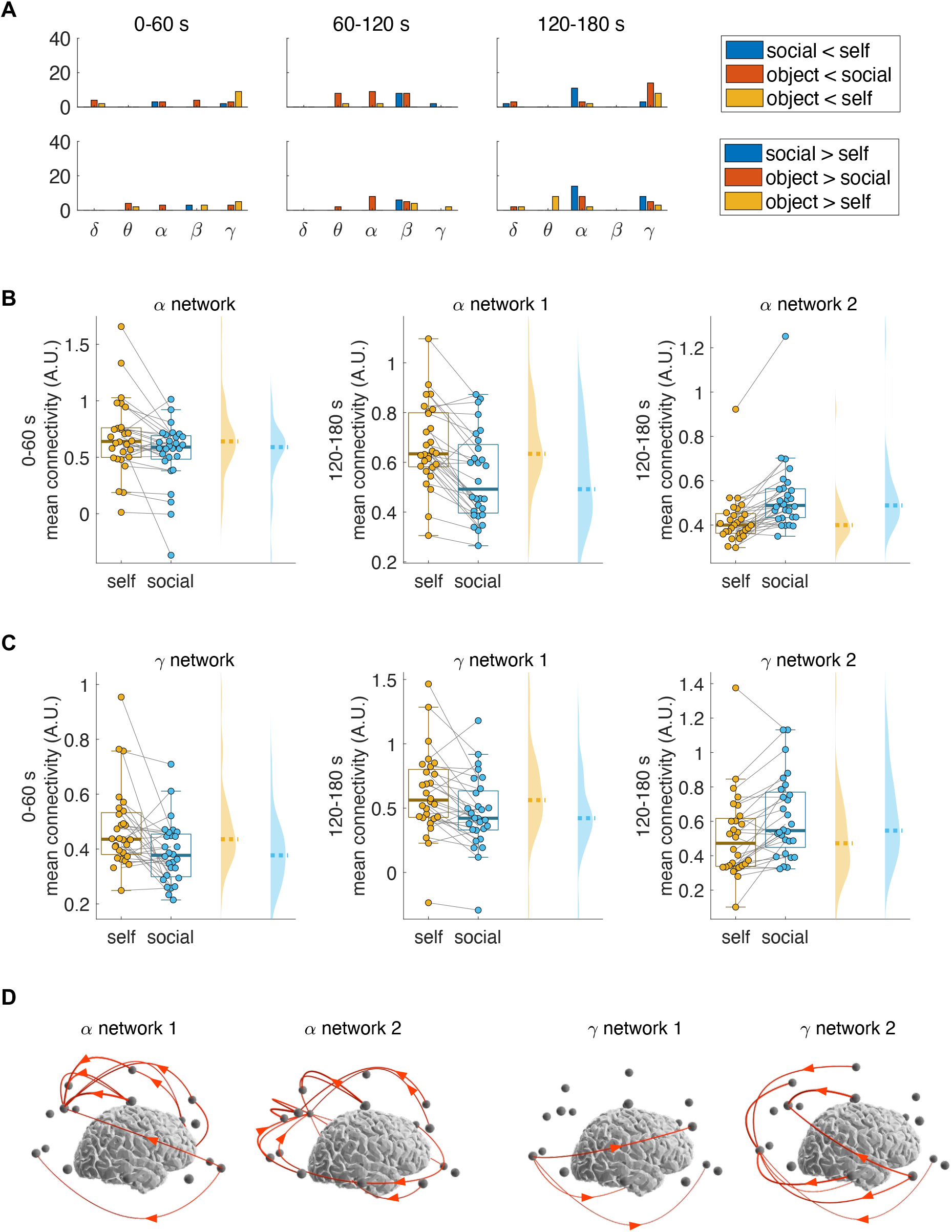
Control analysis on EEG connectivity. (A) Results on the identification of brain networks whose connectivity amplitude differentiate between modalities of touch. The comparisons encompass social vs. self-touch, social vs. object-touch, and self-vs. object touch. The statistical analyses are based on the cluster permutation test to identify the brain networks that showed an increased or decreased amplitude with respect to the touch modalities compared, as specified in the legend. The histograms display the network size, which indicate the number of EEG connectivity links that presented a significant change in their amplitude, between the two touch modalities compared. (B-C) EEG connectivity amplitude change distinguishing social vs self-touch. The main distinctions were found in the EEG connectivity in alpha and gamma brain networks (0-60 s and 120-180s). Data points display the average EEG connectivity amplitude across all links pertaining to the identified network. Each data point corresponds to one participant. (D) Alpha and gamma networks distinguishing self-vs. social touch at 120-180 s. The arrows indicate the directed EEG connectivity links pertaining to the identified network.

Finally, we controlled for differences in brain-heart coupling between self-touch and object-touch. As shown in Figure 2, we observed distinctions primarily in parasympathetic activity, which was higher in later stages, especially in the parasympathetic-theta coupling. The regions involved in these differences spanned a broad frequency range and mainly included short-range connections emerging from central electrodes (see Supplementary material, Figures S1 and S2). In contrast, self-social touch distinctions involved longer-range, bidirectional frontoparietal connections. These findings suggest that the differences between self-touch and object-touch stem from the sensory component associated with self-touch, which involves the arm being touched, a factor absent in object-touch.

## Discussion

Social touch has a vital role in healthy human development^51^; however, the neurophysiology of social touch remains scarcely described^7^. Previous experimental evidence has hinted at a connection between self-awareness and interoceptive mechanisms^52–56^. Therefore, our goal was to investigate the distinctions between social and self-touch by examining brain-heart interactions. In our previous research, we discovered that social and self-touch trigger distinct patterns of activity, both on the neural^12,57^ (fMRI and somatosensory evoked potentials) and autonomic activity^31^. In this study, our objective was to uncover connections between brain networks and the previously observed heartbeat dynamics related to touch^31^. To accomplish this, we employed a method to quantify the relationship between cardiac sympathetic and parasympathetic activities and brain connectivity^49^ within a multimodal touch paradigm. We had previously found that social and self-touch produce different autonomic responses across time, and now we found these responses are directly linked to the interaction of various hubs across the cortex, as captured from scalp recordings. Our findings reveal that social touch, as compared with self or object touch, leads to an increase in the coupling between parasympathetic activity and brain frontoparietal connectivity in the alpha and gamma bands during the early phases of the touch. However, during later stages, social touch caused a decrease in the brain-heart coupling while during self-touch it remained relatively higher, specifically between sympathetic activity and brain connectivity across a wide frequency range, indicating that the social component causes a physiological modulation involving both global brain dynamics and cardiac activity^58^.

Because self-touch probably involves a specific prediction-model underlying sensory attenuation^12,38–40^, we controlled for brain-heart coupling differences between self-object touch. We found that differences occurred in short-range scalp connections emerging from central electrodes, contrasting to self-social touch distinctions involving a longer-range, bidirectional frontoparietal connections. These findings suggest that the differences between self-touch and object-touch stem from primary somatosensory processing^12^, and differences between self-social touch stem from higher order frontoparietal processing^34–37^. Notably, significant differences emerge in the later stages of the touch modalities. We cannot rule out the possibility that these differences may be due to factors such as unpleasantness, boredom, or reduced engagement in the task. This is especially relevant given that parasympathetic-theta coupling changes abruptly at these later stages, potentially indicating variations in arousal^59,60^.

The physiological mechanisms associated with affective touch involve the activation of the tactile receptors located in the skin that are specifically sensitive to light pressure^61^. These signals are transmitted to the C-fibers to ultimately reach the brain through spinal pathways^33,62^. The role of parasympathetic/vagal activity remains poorly understood, although many contexts revealed that affective touch activates the parasympathetic nervous system^63–67^. In our recent research, we showed that touch causes a decrease in cardiac sympathetic activity and an increase in cardiac parasympathetic activity, which was more pronounced in the case of social touch^31^.

We observed that parasympathetic activity increased its coupling with alpha, beta, and gamma oscillations under social touch. Prior studies have associated a variety of brain oscillations in the theta-beta range with affective touch, although these associations were characterized by limited statistical power^20^. Furthermore, these studies revealed distinctions in anterior, central, and posterior scalp regions when comparing affective touch to non-affective touch or to periods of rest ^48^. Touch may not necessarily trigger an increase in a certain EEG power band, but it could rather modulate the communication between different neural hubs^26,27^ in the process of signaling from to the periphery (skin) to the cortex. Indeed, touch processing involves several brain regions including somatosensory, orbitofrontal and cingulate cortices, and the putamen, but also the connectivity between posterior insula with middle cingulate and striatal regions^68–70^. While EEG connectivity patterns may reveal some distinctions in the experimental conditions, our brain-heart coupling analysis provides a more comprehensive view by showing that changes in brain and cardiac activity occur in coordination, rather than independently. However, our analysis does not allow us to identify the exact driver behind this brain-heart coupling. Therefore, we cannot rule out the possibility that a third component may be mediating or moderating this relationship. Still, it is worth noting that our methodology quantifies the coordination of brain-heart dynamics, capturing changes that go beyond simple increases or decreases in brain or cardiac activity. Specifically, our approach uses an information theory-based measure that focuses on the degree of co-fluctuations between EEG and HRV, regardless of their individual amplitudes.

The fact that social touch triggers a distinct pattern of neural activation compared to self-touch suggests that social touch fosters specific mechanisms, unraveled in this study in fronto-parietal networks. These findings concurred cardiac couplings with alpha oscillations predominantly directed from frontal to parietal regions and gamma oscillations from parietal to frontal regions. This distinction may stem from the complex interplay between sensory, emotional, and cognitive processes^34,35,71^, triggered by the social component. Indeed, social touch engages higher-order cognitive functions related to social cognition, empathy, and perspective-taking^58^. The increased cardiac couplings with fronto-parietal networks during social touch may reflect the integration of sensory information with cognitive and emotional appraisal, facilitating the processing of interpersonal dynamics^7^. On the other hand, self-touch, while still eliciting significant physiological responses, may primarily engage sensory and somatosensory processing^44^ without the additional cognitive and emotional components of social interactions. Thus, the neural signatures of self-touch may be characterized by a different pattern of activation, with less impact on cognitive and emotion-related networks. These findings underscore the multifaceted nature of touch perception and its differential effects on brain activity depending on the context in which it occurs.

The brain’s responses to affective touch may be directly associated to the mechanisms of emotion regulation^20^. There is a well-established connection between the brain and the heart in affective situations: numerous studies have examined how emotions influence heart rate and heart rate variability^72–74^, while others have identified links between heart rate fluctuations and specific brain structures^75,76^. These brain-heart interactions are associated with factors such as intensity^59^ and perspective^77^ in humans. Especially, parasympathetic variations are linked to fluctuations in attention, emotional processing^78^, and social engagement^79^. Beyond its impact on emotional processing, the connection between the brain and the rest of the body has been shown to affect body ownership, perspective-taking, and consciousness^42–47^. Considering these connections, it is reasonable to suggest that interoceptive dynamics play a role in the neural mechanisms responsible for distinguishing between social touch and self-initiated touch in the brain. Moreover, exploring brain-heart interactions during social touch can offer deeper insights into the neural mechanisms underlying stress and the stress-buffering effects observed in couple interactions^64,80,81^.

Research into embodiment suggests that humans highly rely on somatosensory inputs^1^, while at the same time multisensory integration is closely associated with interoceptive mechanisms that occur in parallel^47^. For instance, somatosensory detection and tactile action are coupled with the muscle contraction phase of the cardiac cycle and the associated neural responses to heartbeats^82–86^. Cardiac interoception correlates with greater perceived self-body closeness, indicating a link between anchoring the self to the body and improved cardiac interoception^87^. The external manipulation of self-identification through illusions may include disrupted somatosensory perception as well^88^. The posterior insula, which is responsible for interoceptive processing, plays a role in differentiating between the observation of others’ somatosensory experiences and one’s own somatosensory experiences^30^. Higher levels of interoceptive awareness have been associated with stronger effects of social touch representation^89^. The neural processing of cardiac signals in the posterior cingulate cortex was related to bodily self-consciousness, as evidenced by the transient modulations of neural responses to heartbeats that correspond to changes in bodily self-consciousness induced by a full-body illusion^55^. Moreover, neural responses to heartbeats are linked to the self-relatedness of thoughts^52–54^, highlighting the connection between selfhood and the neural monitoring of cardiac inputs.

Our study has limitations, including the use of low-density EEG. We avoided connectivity analysis on source-reconstructed data due to the limited density of scalp recordings, which can lead to biased estimates^90^. Instead, we used a sensor-level approach, which reduces inaccuracies from volume conduction^49,91^. While this method offers valuable insights into connectivity dynamics, caution is needed when interpreting spatial details. Future directions of this research should consider the analysis of functional neuroimaging or high-density source-reconstructed data that address potential volume conduction effects^49,92^.

We identified four limitations in our study related to the specificity of the experimental conditions. First, differences may arise between receiving touch and performing touch. Second, specifically to self-touch, we cannot untangle the potential mechanisms potentially involved in somatosensory processing from the touched arm and those underlying the predictive model evoking self-touch attenuation^12,38–40^. Third, the accuracy of performing the experimental conditions may also play a role. Participants were instructed to perform self- and object touch at an optimal stroking speed for activating C-tactile fibers^18^, but as is the case for all experiments using naturalistic stimuli, there will be a larger degree of natural variation. While this reduces the controllability of stimuli, it increases ecological validity^12,57^ and thereby generalizability to real life conditions compared to within-lab situations using highly controlled stimuli. Finally, tracking the pleasantness over time has proved challenging, as participants may habituate and perceive the touch conditions after the first minute differently. To address this, we conducted separate 1-minute analyses to account for dynamic changes. However, the complexities of physiological changes, especially later in each condition, require cautious interpretation. Specifically, the differences between self-touch and object touch could reflect variations in arousal, unpleasantness, boredom, or reduced engagement rather than purely physiological differences related to self-object touch distinctions.

While we did not delve into examining the directed communication between the brain and heartbeat dynamics or the potential hierarchy of neural avalanches across the cortex and cardiac activity, our framework may have effectively captured feedback and feedforward mechanisms. This aligns with the mechanisms of predictive coding frameworks, which emphasize the integration of interoceptive inputs to anticipate ongoing stimuli^93,94^. Although these insights could deepen our understanding of the physiological mechanisms of affective touch, further investigations are needed to gain a more comprehensive view. This includes studying pathological conditions and the effects of neuromodulation on these mechanisms.

Further advancements in physiological modeling may help us uncover the intricate network structures of interactions across multiple bodily systems. This exploration can shed light on the complex and hierarchical organizations that come into play during various physiological and cognitive states, including experiences related to affective touch. Various interactions between the brain and other organs have been documented during different conscious experiences^47^. Using computational approaches that account for the numerous mechanisms these interactions manifest within brain-other organ systems—whether they are coupled, intertwined, or integrated—can provide valuable insights into the physiological foundations of our overall conscious experience.

A deeper comprehension of the neurophysiology of touch holds significant relevance. It extends from its influence on the sense of body ownership, as evidenced by its role in enhancing this sense, as seen in the correlation between touch pleasantness and the degree of subjective embodiment in the rubber hand illusion^41,95^. Moreover, this understanding has critical clinical implications^96^, particularly in the context of treating disorders related to disrupted pleasant and unpleasant touch processing, which is observed in certain conditions, including autism^97^, anorexia nervosa^98^, and schizophrenia^99^. Understanding the fundamental mechanisms that guide touch pathways is essential, as disruptions in these pathways can result in changes to the sensitivity and specificity of tactile receptors, resulting in the perception of tactile stimuli as uncomfortable or even painful^61^.

By advancing our understanding of the large-scale neural interactions linked to social touch, we can gain valuable insights applicable to the treating pathological conditions. This includes cases of pathological painful touch, where such insights could complement, for instance, markers based on event-related potential analysis^100^. At a fundamental level, this research could establish potential connections with the mechano-sensation mechanisms at cellular level linking brain, skin and heart^101–103^. This, in turn, could potentially lead to the development of strategies for mitigating pain through interventions that trigger brain-heart pathways.

## Conclusion

We revealed a connection between touch, particularly social touch, and the interaction of cardiac and cortical dynamics. Our results hold potential clinical relevance, offering insights into the neurophysiology of touch, particularly in the investigation of conditions where a disrupted touch processing is found.

## Materials and Methods

### Participants

This is a retrospective analysis of a cohort undergoing a multimodal touch paradigm^31^. A total of 28 healthy adult volunteers participated in this study (16 females, mean age 29.04 years, SD=5.16). Participants were required to be fluent in English, and had no current cardiac, sensory/motor, or affective/psychiatric conditions. Data acquisition was performed at the Center for Social and Affective Neuroscience (CSAN), Linköping, Sweden. All participants provided informed consent in accordance with the Helsinki Declaration and were compensated for their participation. The study was approved by the Swedish ethics board.

The participants engaged in an established experimental task known as the self-other-touch paradigm. The task employed a randomized block design and encompassed three distinct conditions: social touch (being stroked on the left forearm by the experimenter), self-touch (stroking of the participant’s own left forearm), and object-touch (participant stroking a pillow). Each of the three conditions lasted 180 seconds. For the self and object-touch condition, participants were instructed to gently stroke, mimicking the touch they would use when interacting with someone they like, using their right hand.

Instructions for each block of the task were presented on a screen. The instructions, provided in English, were displayed including the following prompts: “Social touch: Your arm will be touched by the experimenter», «Self-touch: Please stroke your arm”; “Object-touch: Please stroke the object”; Participants either received stimulation or performed the stimulation themselves, continuing for a duration of 3 minutes. Participants performed the task with their eyes closed, therefore the experimenter informed them when the condition was over. The female experimenter (PS) stationed adjacent to the participant, but out of the peripheral view (blocked by a desk divider) replicated the participant’s movements in the same area of the forearm as closely as possible.

Touch was instructed to be performed at optimal stroking speed, which would activate C-tactile fibers and result in a sensation typically described as pleasant^104^. This type of touch is often referred to as affective touch in the literature^17,18^.

The EEG and ECG recordings were obtained using *B-Alert* with 20 channels and BIOPAC MP160 for additional channels for triggers. ECGs were recorded using additional 2 electrodes dedicated to record ECG from the chest. Recordings were performed with *Acqknowledge* software (*Biopac*) at 2000 Hz using mastoids as references.

### EEG and ECG data processing

EEG data were acquired using a 20-channel BioSemi ActiveTwo system, together with a one-lead ECG, sampled at 512 Hz. During data collection, the participants were seated comfortably and with eyes closed.

The EEG and ECG data were pre-processing using MATLAB R2022b and Fieldtrip Toolbox^105^. The EEG and ECG data were bandpass filtered with a Butterworth filter of order 4 between 0.5 and 45 Hz. Large movement artifacts were removed from EEG using a wavelet-enhanced independent component analysis^79^. Independent Component Analysis (ICA) was then re-run to detect and set to zero the components with eye movements and cardiac-field artifacts. ECG was included in the ICA computation to improve the process of identifying cardiac artifacts. EEG channels were re-referenced using a common average^106^.

The R-peaks from the ECG were identified using an automatized process, followed by an automated inspection of misdetections and manual correction if required. The procedure was based on a template-based method for detecting R-peaks. For the correction of misdetection, all the detected peaks were visually inspected over the original ECG, along with the marks on potentially misdetected heartbeats and the inter-beat intervals histogram.

### Computation of brain-heart interactions

Brain-heart interactions were computed using a framework that quantifies the coupling between brain connectivity and cardiac sympathetic or parasympathetic indices^49^.

The estimation of cardiac sympathetic and parasympathetic activities was based on a method that uses the fluctuating geometry of the Poincaré plot constructed from inter beat intervals (IBI)^50^. The method combines the time-resolved quantification of the baseline cardiac cycle duration (CCD), the short-term (SD_1_) and long-term (SD_2_) fluctuations of heart rate variability. The fluctuations of the Poincare plot-derived measures were computed with a sliding-time window, as shown in Eq. 1, 2 and 3:

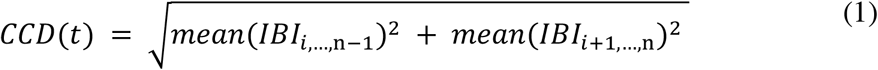

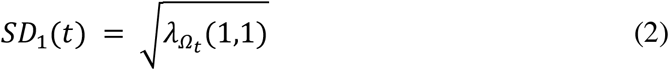

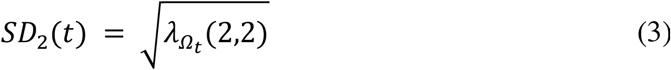

where 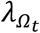 is the matrix with the eigenvalues of the covariance matrix of *IBI*_i,…,n…1_ and *IBI*_i+1,…,n_, with *Ω*_*t*_: *t* – *T* ≤ *t*_i_ ≤ *t*, and *n* is the length of IBI in the time window *Ω*_*t*_. In this study T is fixed in 15 seconds, as per previous simulation studies in humans ^81^. The distance to the origin *CCD*_0_ and ellipse ratios *SD*_01_ and *SD*_02_ for the whole experimental duration are computed to re-center the time-resolved estimations of CCD, SD_1_ and SD_2_. Then, the Cardiac Parasympathetic Index (*CPI*) and the Cardiac Sympathetic Index (*CSI*), are computed as follows:

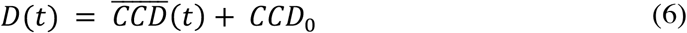

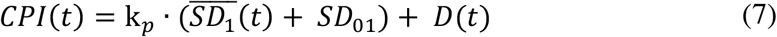

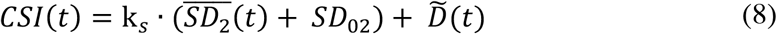

where 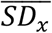 is the demeaned *SD*_*x*_ and 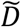 is the flipped *D* with respect the mean. The coefficients k_*p*_ and k_*s*_ define the weight of the fast and slow HRV oscillations, with respect the changes in the baseline heart rate. In this study, the values were defined as k_*p*_ = 10 and k_*s*_ = 1. Those values were chosen based on the well-stablished effects of autonomic modulations on cardiac dynamics: Sympathetic modulations primarily influence baseline heart rate, but also slower HRV changes, while parasympathetic modulations are typically captured by quantifying faster HRV changes^50^.

The EEG spectrogram was computed using the short-time Fourier transform with a Hanning taper. Calculations were performed through a sliding time window of 2 seconds with a 50% overlap, resulting in a spectrogram resolution of 1 second and 0.5 Hz. Time series were integrated within five frequency bands (delta: 1-4 Hz, theta: 4-8 Hz, alpha: 8-12 Hz, beta: 12-30 Hz, gamma: 30-45 Hz). The directed time-varying connectivity between two EEG channels was quantified using an adaptative Markov process, as shown in Equation *(7)*, where *f* is the main frequency, *θ*_*f*_ is the phase (*f = 1*, …, *45 Hz)*. The model estimates the directed connectivity at a specific frequency band (*F =* {*delta, theta, alpha, beta, gamma*}) using least squares in a first order auto-regressive process with an external term, as shown in *(8)*, where *A*_*F*_ is a constant and *ε*_*F*_ is the adjusted error. Therefore, the directed connectivity is obtained from the adjusted coefficient from the external term *B*_*F*_, as shown in Equation *(9)*.

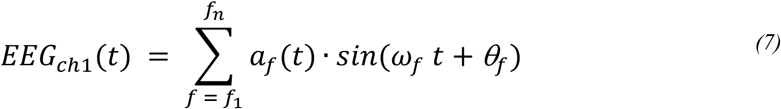

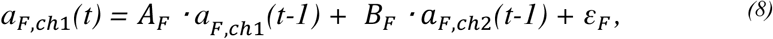

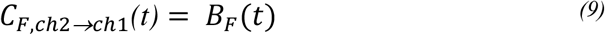

As depicted in Figure 1, brain-heart coupling was quantified by considering the relationships between brain connectivity fluctuations and cardiac sympathetic-parasympathetic indices. The brain-heart coupling was assessed using Maximal Information Coefficient (MIC). MIC is a method that quantifies the coupling between two time series^107^. MIC evaluates similarities between different segments separately at an adapted time scale that maximizes the mutual information, with a final measure that wraps the similarities across the whole time-course. The Equations *(10)* and *(11)* show the MIC computation between two time series X and Y. The mutual information *I*_g_ is computed to different grid combinations *g* ∈ *G*_*xy*_. The mutual information values are normalized by the minimum joint entropy log_2_ min{*n*_*x*_, *n*_*y*_}, resulting in an index in the range 0-1. Then, the quantified coupling between X and Y corresponds to the normalized mutual information resulting from the grid that maximizes the MIC value.

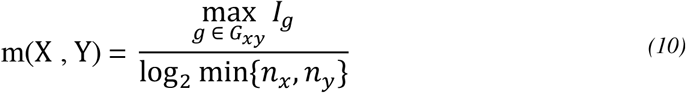

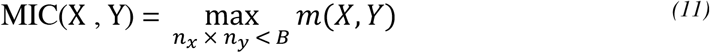

where B = *N*^0.6^, and N is the dimension of the signals^107^.

Finally, one MIC value is obtained for all possible combinations of frequency band, directed connectivity for all pair of EEG channels and their coupling to either cardiac sympathetic or parasympathetic activity.

### Statistical analysis

MIC values were compared between experimental conditions: social touch (being stroked on the forearm by the experimenter), self-touch (stroking of the participant’s own forearm), and object-touch (participant stroking a pillow). Each of the three conditions lasted 180 seconds, that were analyzed in the segment 0-60, 60-120 and 120-180 seconds. Statistical comparisons were based on two-sided Wilcoxon signed rank tests for pared comparisons. P-values significance was evaluated by using cluster-permutation analyses. Clustered effects were revealed using a non-parametric version of cluster permutation analysis^108^. Cluster permutation analysis was applied to the MIC values computed between the directed connectivity and the cardiac sympathetic/parasympathetic activity separately. Cluster-based permutation test included a preliminary mask definition, identification of candidate clusters, and the computation of cluster statistics with Monte Carlo’s p-value correction. First, the preliminary mask was defined through Wilcoxon test, with alpha = 0.05, to the 992 MIC values corresponding to all the possible pairs of channel combinations in both directions. The identification of neighboring points was based on neighborhood definition for the 20 EEG channels. A minimum cluster size of 3 neighbors was imposed. Cluster statistics were computed from 10,000 random partitions. The proportion of random partitions that resulted in a lower p-value than the observed one was considered as the Monte Carlo p-value, with significance at alpha = 0.05. The cluster statistic considered is the Wilcoxon’s absolute maximum Z-value obtained from all the samples of the identified networks, separately.

The visualization of the brain networks coupled with heartbeat dynamics was performed using Vizaj^109^. Distributions of the mean brain-heart coupling across the identified networks are displayed as individual data points together with their estimated distributions and box plots. Box plots display the median value, the edges indicate the interquartile range (IQR), and the whiskers extend from the edges within 1.5 times the IQR from the respective quartiles.

## Data availability

Data are available upon reasonable request to the corresponding author.

## Code availability

Codes are available at https://github.com/diegocandiar/heart_brain_conn, https://github.com/diegocandiar/robust_hrv, and https://github.com/diegocandiar/eeg_cluster_wilcoxon.

## Author contributions

Diego Candia-Rivera: Methodology, Visualization, Formal analysis, Writing – first version of the manuscript, Writing – review & editing.

Fabrizio de Vico Fallani: Visualization, Formal analysis, Writing – review & editing.

Rebecca Boehme: Experimental design, Formal analysis, Writing – review & editing. Paula C. Salamone: Experimental design, Data curation, Formal analysis, Writing – review & editing.

## Competing interests

Nothing to declare.

## Acknowledgements

This research was supported by the Fredrik och Ingrid Thurings stiftelse, the Lions Forskningsfond and the Swedish Research Council (2019-01873).

## Notes

### Competing Interest Statement

The authors have declared no competing interest.

### Summary of Updates

New results and figures included after 2 rounds of revision

